# Hydrogel encapsulation of a designed fluorescent protein biosensor for continuous measurements of sub-100 nanomolar nicotine

**DOI:** 10.1101/2024.12.02.625538

**Authors:** Aaron L. Nichols, Christopher B. Marotta, Daniel A. Wagenaar, Stephen L. Mayo, Dennis A. Dougherty, Henry A. Lester

## Abstract

The reinforcing and addictive properties of nicotine result from concentration- and time-dependent activation, desensitization, and upregulation of nicotinic acetylcholine receptors. However, time-resolved [nicotine] measurement in people who consume nicotine is challenging, as current approaches are expensive, invasive, tedious, and discontinuous. To address the challenge of continuous nicotine monitoring in human biofluids, we report the encapsulation of a purified, previously developed fluorescent biosensor protein, iNicSnFR12, into acrylamide hydrogels and polyethylene glycol diacrylate (PEGDA) hydrogels. We optimized the hydrogels for optical clarity and straightforward slicing. With fluorescence photometry of the hydrogels in a microscope and an integrated miniscope, [nicotine] is detected within a few min at the smoking- and vaping-relevant level of 10 - 100 nM (1.62 – 16.2 ng/ml), even in a 250 µm thick hydrogel at the end of 400 µm dia multimode fiber optic. Concentration-response relations are consistent with previous measurements on isolated iNicSnFR12. Leaching of iNicSnFR12 from the hydrogel and inactivation of iNicSnFR12 are minimal for several days, and nicotine can be detected for at least 10 months after casting. This work provides the molecular, photophysical, and mechanical bases for personal, wearable continuous [nicotine] monitoring, with straightforward extensions to existing, homologous “iDrugSnFR” proteins for other abused and prescribed drugs.

---

A billion persons smoke nicotine daily, and an estimated 60% would like to quit. It has long been appreciated that the time course of [nicotine] near nicotine acetylcholine receptors (nAChRs), which we term *[nicotine]*_*t*_, partially determines both nicotine addiction and successful nicotine replacement therapy. ^1-5^

A wearable continuous nicotine monitor would serve in research on four topics that involve measuring *[nicotine]*_*t*_. (1) Researchers wish to “know the enemy”, the ways in which *[nicotine]*_*t*_ varies among individuals who smoke, vape, and use oral nicotine products. (2) It will also be necessary to “know the therapy”. FDA-approved nicotine replacement therapy (NRT) may soon be enhanced by trials of prescription mesh nebulizers; and it is necessary to study how *[nicotine]*_*t*_ correlates with success in smoking cessation. (3) The research community would like to “know the physiology”: by measuring *[nicotine]*_*t*_, researchers can correlate the contributions of activation, desensitization, and upregulation of nAChRs to reinforcement, withdrawal, and addiction. (4) A wearable continuous nicotine monitor will produce data during *ad libitum* nicotine ingestion.

Genetically encoded fluorescent biosensors present one potential strategy for the generation of a continuous nicotine monitor in animals and reduced systems. To date, biosensors have been used to study endogenous molecules such as serotonin, GABA, glutamate, and dopamine, as well as abused and prescribed drugs – including nicotinic agonists, opioids, rapidly acting anti-depressants, and selective-serotonin reuptake inhibitors (SSRIs). ^6-16^ Genetically encoded biosensors have been utilized in mammalian cell culture, primary neuronal culture, and model organisms, via transfection and viral transduction.

Related techniques for gene transfer cannot be used in humans for pharmacokinetic monitoring of drugs. Instead, an approachable tactic may be to build a continuous fluorescent monitor by entrapping a soluble fluorescent biosensor protein into a transparent hydrogel. Hydrogels have been extensively studied for tissue engineering, biomimetics, biocatalytic scaffolds, and even a continuous glucose monitor (CGM).^17-19^ Non-degrading hydrogels provide excellent parameters for a medical device – providing an aqueous environment that allows small molecule diffusion into a network, while simultaneously excluding larger biomolecule diffusion. A successful hydrogel network restricts off-target binding, limits chemical interference, and reduces protein degradation, while also slowing diffusion of an introduced protein out of the hydrogel.^20^ Encapsulating target proteins during hydrogel formation allows for the design of needed structures through mold casting. In addition, hydrogel synthesis utilizing photopolymerization allows 3D printing of the desired structures for optimal device designs.^21^ This flexibility allows device construction around the excitation and emission needed for fluorescence detection.

Dermal interstitial fluid (ISF) can be probed with minimally invasive techniques. ISF *[nicotine]*_*t*_, is expected to closely resemble *[nicotine]*_*t*_, in blood and in cerebrospinal fluid ^3, 22^. The highest [nicotine] in these three compartments occurs during the bolus of nicotine from puffing on a cigarette/vape, ∼ 100 nM. This phase largely governs the reinforcing effects of nicotine.^23^ When the smoking / vaping session ends, *[nicotine]*_*t*_ partially declines within a few minutes Then a prolonged, declining phase begins, with a time constant of 2 – 4 h. This tapering *[nicotine]*_*t*_ suppresses withdrawal symptoms and maintains the cellular biological aspects of addiction.^24^ While we know the average *[nicotine]*_*t*_ for various modes of nicotine ingestion, individual variability in nicotine processing is considerable.^23, 25, 26^

Two methods currently provide “gold standard” measurements of *[nicotine]*_*t*_. 1) Intravenous (IV) blood draws provide instantaneous time point measurements of nicotine concentration in the blood, but each IV sample is costly and tedious to execute and process.^27^ 2) Positron emission tomography has better time resolution (a few s) but also comes with a high reagent and machine cost. ^28^

While some indirect, proxy measures for estimating nicotine concentration in biofluids exist, such as the nicotine metabolite ratio (NMR), these measurements cannot directly capture *[nicotine]*_*t*_.^29, 30^ Electrochemical nicotine sensors based on nicotine-oxidizing enzymes lack the molecular amplification of fluorescence and are, therefore, 20 - 100-fold less sensitive than fluorescence. ^31-33^ To date, electrochemical measurements have been tested only on sweat.^31-33^ which is less desirable than interstitial fluid because of inherent time delays and acid trapping of nicotine. ^34^

We report *in vitro* continuous fluorescence measurements of *[nicotine]*_*t*_ with the most evolved nicotine biosensor, iNicSnFR12,^35^ encapsulated in a hydrogel. We adapted techniques from measurements using genetically encoded fluorescent biosensors on intact rodents (a miniature integrated fluorescence microscope, fiber photometry) and brain slices (vibratome slicing, immobilization by a harp). This work provides a first step toward the employment of designed fluorescent biosensors for personalized monitoring of *[nicotine]*_*t*_ in human ISF.

## Experimental

### Hydrogel formation

#### Acrylamide hydrogels

A 500 mM, pH 7.5 solution of TEMED was generated for acrylamide hydrogel reactions. 4 mmol N,N dimethylacrylamide monomer was crosslinked with 1 mmol tetra(ethylene glycol) diacrylate using a 0.025 molar ratio of TEMED as a co-initiator and 0.0025 molar ratio ammonium persulfate (APS) as a radical initiator, in the presence of 0.2x PBS, pH 7.4 and 0.5–10 μM final concentration of biosensor protein.

To ensure proper mixing and stability of biosensor protein stepwise addition of the components to a glass vial was as follows: Deionized (DI) water, monomer, crosslinker, TEMED, 1x PBS, pH 7.4, and biosensor protein. The vial was sealed with a rubber stopper and the solution was sparged with argon for ∼30 s. The vial was cooled in an ice bath for 5 min, the rubber stopper was removed, the APS was rapidly added, the rubber stopper was replaced, the vial was gently swirled and placed immediately back in the ice bath for 1-2 h. The vial was then held at 4 °C for 1-2 h. Next, the vial was uncapped, and the formed hydrogel was rinsed with DI water. The vial was cracked to remove the hydrogel puck. The puck was washed twice using 50 mL of 1X PBS, pH 7.4 in a 50 mL conical tube for 1-2 h. The puck was then washed overnight in 50 mL of 1X PBS, pH 7.4.

#### PEGDA/ Irgacure hydrogels

A 100 mg/mL stock of Irgacure 2959 (2-hydroxy-4′-(2-hydroxyethoxy)-2-methylpropiophenone)) was solvated in ethanol/water as a 70% v/v mix. Poly(ethylene glycol) diacrylate (PEGDA) (M_n_ 700) and Irgacure 2959 stock were mixed in 1X PBS, pH 7.4 at 46% w/w and 4.4% w/w. The resulting mixture was combined in a 1:1 ratio with 20 μM iNicSnFR12 solution to a final concentration of 23% PEGDA, 2.2% Irgacure 2959, 10 μM iNicSnFR12 in 0.97X PBS, pH 7.4, 3% ethanol.^21^ The mixture was irradiated on ice at 100% power using a 365 nm, 1350 mW LED, 1700 mA for 10 mins. The hydrogel puck was washed twice in 50 mL of 1X PBS, pH 7.4 in a 50 mL conical tube for 1-2 h. The puck was then washed overnight in 50 mL of 1X PBS, pH 7.4.

The addition of laponite (2.5%)^21^, rat collagen type I (50%), hyaluronic acid (4%), or bovine gelatin (5%) were tested as thickening agents for PEGDIrgacure formation. These agents gave suboptimal results, including mottled translucence and insufficiently hard hydrogels, and were consequently abandoned.

#### Hydrogel leaching tests

To test leaching of the biosensor from acrylamide and PEGDA/Irgacure hydrogels, the 50 mL wash was concentrated to ∼300 μL and measured on a Spark M20 96-well fluorescence plate reader (Tecan), with excitation at 485 nm and emission at 535 nm. Only background fluorescence was observed

### Epifluorescence imaging of hydrogels

Acrylamide hydrogels could be manually chipped to uneven slices at 0.5-1.0 μm thick. PEGDA-Irgacure hydrogels were sliced at 250 μm using a Leica VT 1000S Vibratome with a razor blade. Hydrogels were immobilized using manually placed weighted wires and/or “harps” usually intended to immobilize brain slices.

Time-resolved concentration–response imaging was performed as described ^36^ on a modified Olympus IX-81 microscope (wide-field epifluorescence mode using a 10× lens, λ_ex_ = 470 nm λ_em_ = 535). Images were acquired at 1 frame/s with a back-illuminated electron multiplier CCD camera (iXon DU-897, Andor Technology) controlled by Andor IQ3 software.

Solutions were delivered from elevated reservoirs ^36, 37^ by gravity flow via solenoid valves, providing solution changes in the imaging chamber with a time constant < 10 s. The vehicle was 1x PBS, pH 7.4. Data analysis procedures included subtraction of blank (non-hydrogel) areas and corrections for baseline drifts using OriginPro 2018 software. Displayed time series data were smoothed by moving averages over 30 s.

#### Miniscope imaging of hydrogels

Using the open-source instructions, a v4.4 UCLA fluorescent miniscope kit was assembled.^38, 39^ The elevated reservoirs and gravity flow setup described above were adapted so that the miniscope functioned as an inverted fluorescent fiber photometer. We 3-D printed an adapter to hold both the miniscope and the ferrule of a Thorlabs fiber optic (400 μm dia, 0.39 NA, 10 mm length). The adapter was held by a micromanipulator rod that usually holds an electrophysiology headstage. The instrument end of the fiber optic was placed at the focus of the miniscope objective lens, and the entire image of the fiber optic was averaged for the data shown. The near-tissue end of the fiber optic was flush with the bottom of a 160 µm cover slip. Atop the cover slip, we placed a vibratome-sliced hydrogel immobilized with a harp. Images were collected at the slowest available rate for the miniscope, 10 /s. Except for Figure 3B below, displayed time series data were smoothed by moving averages over 30 s.

## Results and Discussion

### Generation of hydrogels compatible with bacterially expressed and purified fluorescent proteins

Our first generation of hydrogels was composed of iNicSnFR12 Q368C (a nicotine biosensor with a point mutation as a chemical handle of iNicSnFR12)^35^ encased in polymerized acrylamide at a final concentration of 2 μM. These hydrogels were brittle and fragile, requiring manual chipping to produce samples for use in wide-field epifluorescence experiments. The acrylamide hydrogel showed an even distribution of biosensor throughout the hydrogel, with no detectable puncta. (**Figure 1A**).

**Figure 1.**
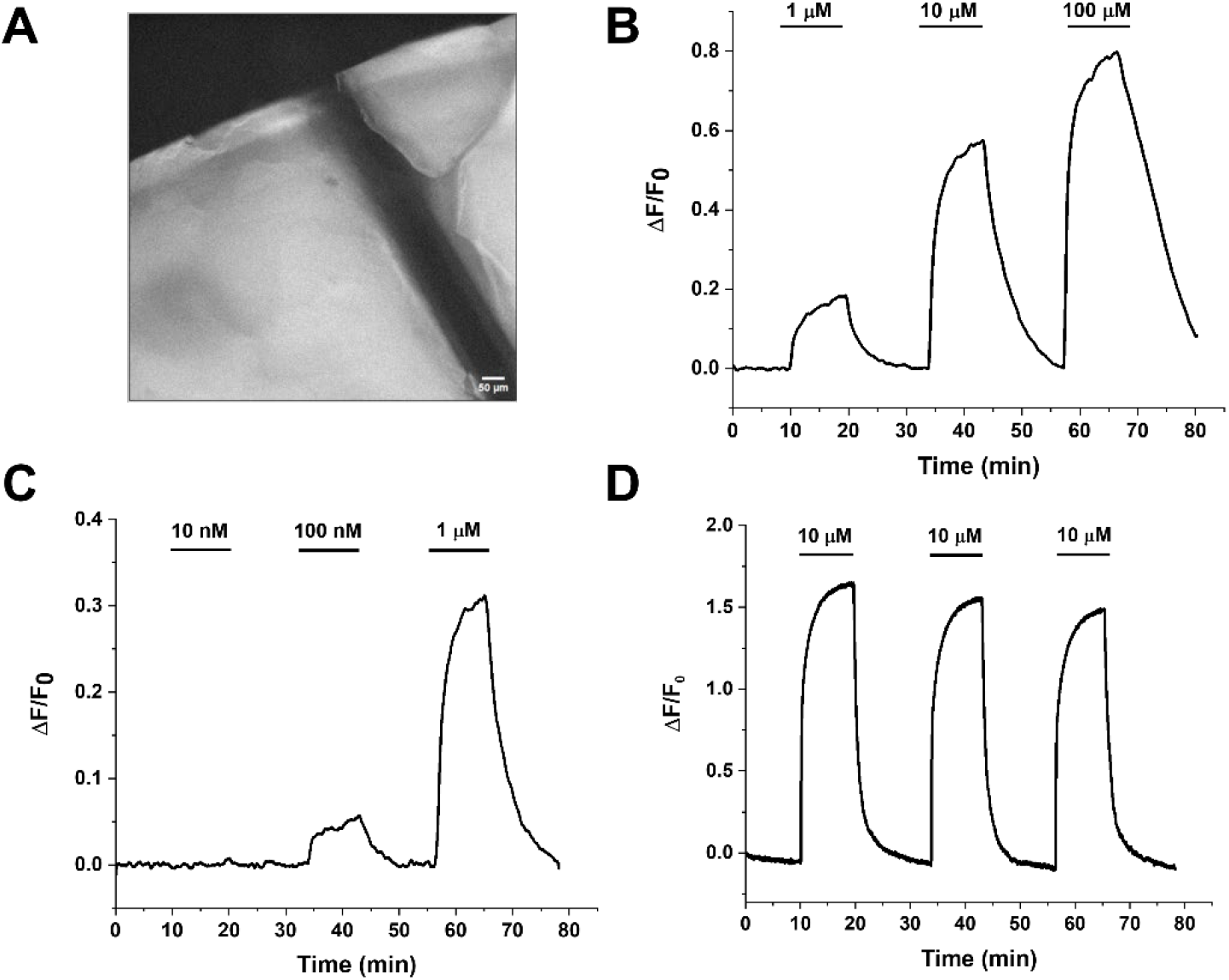
iNicSnFR12-Q368C in acrylamide hydrogels. (A) Widefield image of acrylamide hydrogel. The dark line is a weighted wire. Scale bar, 50 μm. (B, C). Time-resolved concentration-response relations over a [nicotine] range from 100 nM to 100 µM in two exemplar hydrogels. Wash-in and washout kinetics are slower than solution changes; see text for analysis in terms of diffusion. (D) In another gel, repeated applications of the same dose of nicotine show a robust response over 75 min.

We performed time-resolved concentration-response relations with nicotine on the hydrogel slices. We observed a robust fluorescent response to nicotine across a range of [nicotine] ≥ 100 nM (**Figure1B-1C)**. The observed wash-in and washout kinetics characteristics were slower than the solution changes and were several fold slower than previous results with iNicSnFRs in mammalian cell culture and primary mouse hippocampal neurons.^13, 35, 37^ Repeated applications of nicotine to the acrylamide hydrogel showed minimal rundown, with a fluorescent decrease of 7-8% over the course of a 75 min experiment.(**Figure 1D**). For all dose response relations, we observed slower wash-in and washout kinetics for the interior of the hydrogel versus more solvent- exposed regions, suggesting that diffusion limited the access of the nicotine to the sensor protein. (Supporting Information: Analysis of solvent proximity to fluorescent response of iNicSnFR12 acrylamide hydrogel) The fluorescent response of iNcSnFR12 Q368C in concentration-response relations with nicotine indicated functional protein, which demonstrated that a chemical handle would not be needed to stably encase iNicSnFR12 in the hydrogel. Consequently, subsequent hydrogel experiments used iNicSnFR12 itself.

### PEGDA/Irgacure hydrogels

While the initial data for polyacrylamide gels showed that iNicSnFR12 biosensor was functional in a hydrogel, it seemed possible to improve on the formulation in three ways: 1) A less toxic hydrogel matrix than acrylamide would be better suited for use in biological systems; 2) Improved mechanical properties would allow more robust manipulation and handling; and 3) Gel-like consistency would allow systematic shaping and slicing.

To accomplish these goals, we investigated a PEGDA/Irgacure hydrogel formulation.^21^ After experimenting with the addition of thickening agents (including laponite, rat collagen Type 1, hyaluronic acid, and bovine gelatin), we determined that a PEGDA/Irgacure iNicSnFR12-containing hydrogel with no thickening agents provided the best clarity and flexibility.

We cast PEGDA/Irgacure hydrogels at concentrations between 0.1 μM and 50 μM iNicSnFR12. Hydrogels with iNicSnFR12 concentration <10 μM provided less fluorescence than desired for accurate nicotine detection, while increasing [iNicSnFR12] above 10 μM did not markedly increase the detectable signal. Additionally, increasing the biosensor concentration to 50 μM resulted in fluorescent puncta, rather than a homogeneous distribution of fluorescence. (Supporting Information: Widefield imaging of varied PEGDA hydrogel preparations). Therefore, subsequent experiments used 10 μM iNicSnFR12 in the PEGDA/Irgacure hydrogels.

### Vibratome-sliced PEGDA hydrogels

To obtain reproducible hydrogel thickness, we adapted brain slicing techniques to the PEGDA/Irgacure hydrogel: we obtained 250 μm thick slices using a vibrating microtome (“vibratome”) (**Figure 2A)**. Slices generated in this fashion had minimal opacity and exhibited homogeneous distribution of fluorescence (**Figure 2B**). Concentration-response relations with 250 μm thick slices showed a robust fluorescent response of iNicSnFR12 to nicotine across a range of concentrations.

**Figure 2.**
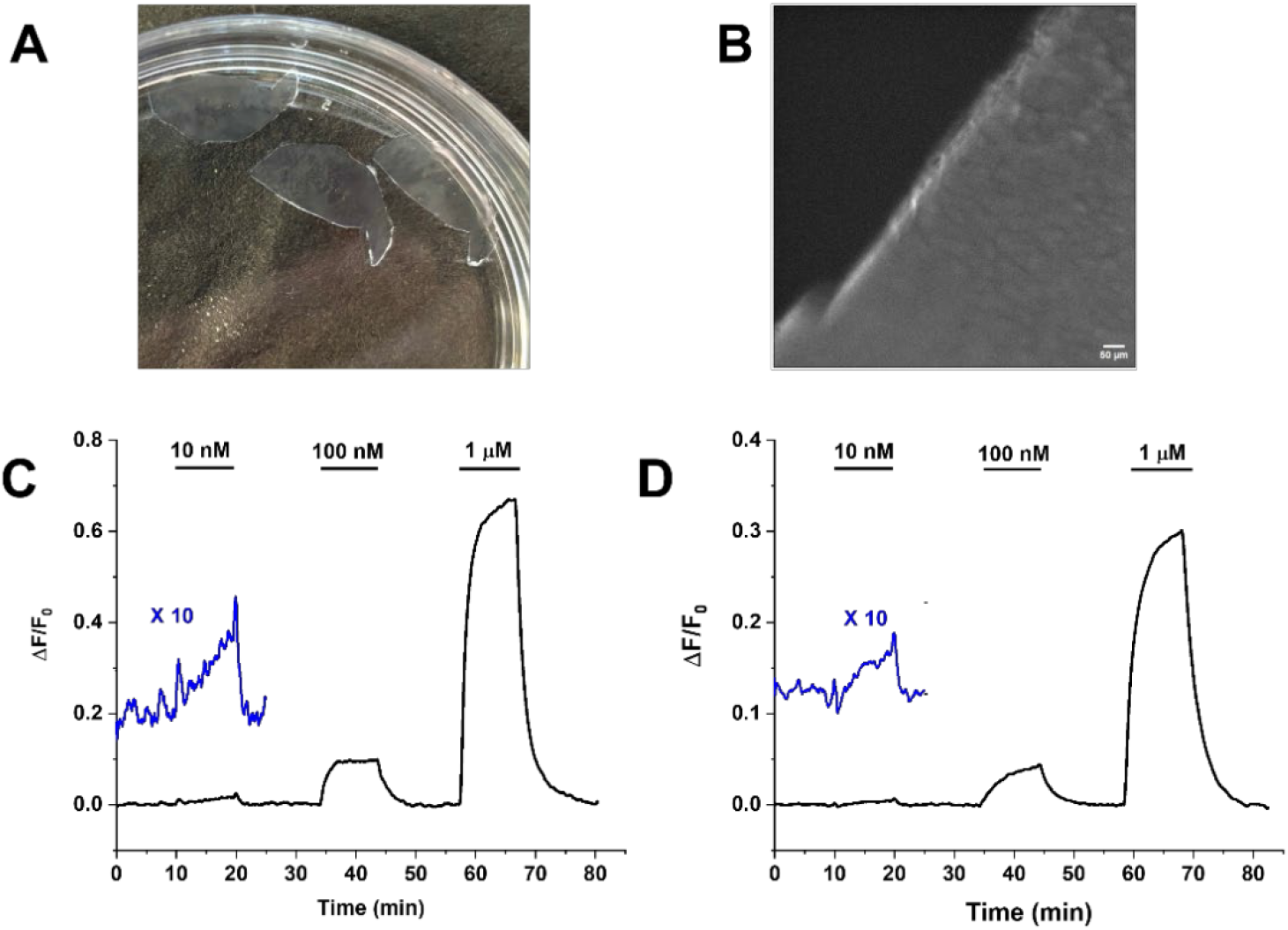
Experiments with iNicSnFR12 PEGDA hydrogels, 250 μm thick. (A) Photograph of hydrogels in a 35 mm dish. Hydrogel slices have optical clarity. (B) Widefield fluorescence image. Scale bar, 50 μm. (C) Time-resolved concentrationresponse relations to nicotine, 10 nM, 100 nM, and 1 μM. Wash-in and washout kinetics are slower than solution changes; see text for analysis. (D) Ten months after casting; storage at 4° C. The iN-icSnFR12 PEGDA hydrogel shows a reduced, but still measurable response, In C and D, the blue trace shows the 10 nM response, magnified 10-fold vertically.

**Figure 3.**
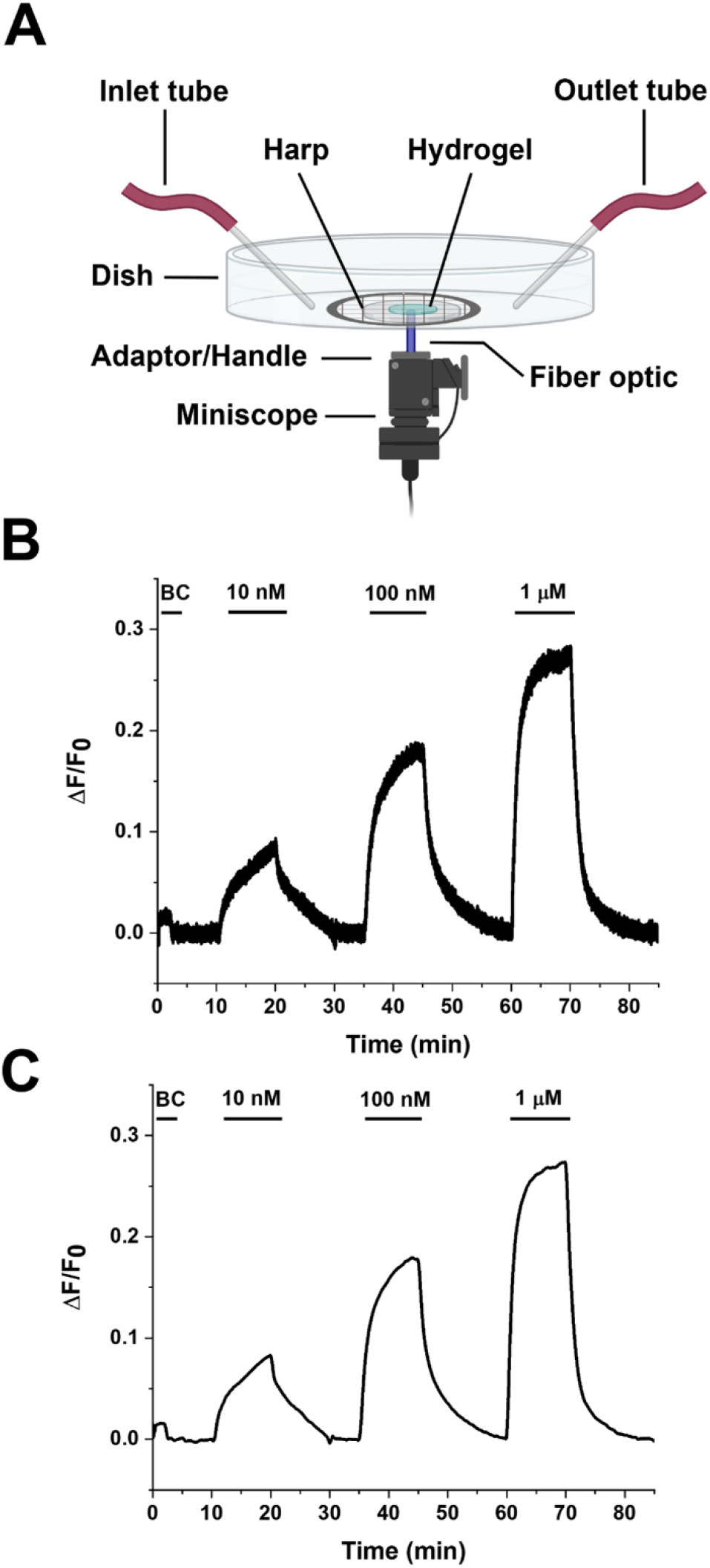
Fiber photometry of iNicSnFR12 PEGDA hydrogel using an integrated miniature fluorescence microscope (miniscope). (A) Schematic. The miniscope is inverted from its usual position on a rodent head. Nicotine solutions are applied through the inlet tube and removed by suction through the outlet tube. A harp immobilizes a 250 μm PEGDA hydrogel slice. (B) iNicSnFR12 PEGDA hydrogel detects [nicotine] as low as 10 nM. BC, zeronicotine buffer control, showing artifact due to solution changes. (C) Smoothed miniscope data using adjacent averaging of 300 frames (30 s).

### Concentration dependence of nicotine-induced fluorescence

In each of four slices tested (examples in **Figure 2C, D**), we observed responses at [nicotine] as low as10 nM. The 10 nM waveforms were noisy and distorted by mechanical artifacts, small changes in local pH, and other idiosyncrasies of the iNicSnFR series noted in previous papers.^13, 15^ Qualitatively, the 10-fold scaled 10 nM waveforms in Figure 3 C,D are comparable in size to the 100 nM waveforms, consistent with the observed Hill coefficient of 1 for the iNicSnFR series. ^13, 15, 35^

The 1 μM responses in **Figures 2 C,D** are 7.1-fold larger than the 100 nM responses, markedly less than the 10-fold of a linear relationship. This suggests that 1 μM is an appreciable fraction of the EC_50_ for the iNicFR12-nicotine dose-response relation. In a previous study, the EC_50_ was 6 – 9 μM; this parameter may vary by several fold with small changes in pH, ionic strength, and temperature,^15^ and perhaps the hydrogel environment.

### Semiquantitative Analysis of diffusion

The rise times (10% - 90%) of the responses at 1 μM nicotine were 5 – 7 min (Figure 2C, D). The experimental chamber changed solutions within 10 s, much faster than these rise and fall times. In stopped-flow measurements, the iNicSnFR series responds to jumps of [nicotine] within 1 s.^13, 15^ Therefore, we present a semiquantitative analysis of the PEGDA hydrogel waveforms based on diffusion properties. ^40^ As expected for a diffusion process, the rise and decay waveforms were the sum of several exponential terms.

We assumed that nicotine undergoes diffusion within the hydrogel plane of uniform thickness *x* = 250 μm. We assumed that [nicotine] at the upper surface of the hydrogel is jumped instantaneously to the level in the perfusate, and the lower surface, at the cover slip, is closed to access from the solution. We also assumed that the free solution diffusion constant for nicotine, *D*, equals 0.5 μm^2^ / ms, typical of low-MW alkaloids.^41^ We assumed that the effective diffusion constant *D*_*eff*_ *= D/ab*, where *a* = 1.5 for diffusion within the restricted space of the hydrogel. The factor *b* represents rebinding to the iNicSnFR molecules and equals K_d_ / ([iNicSnFR12] +K_d)_). Assuming that the K_d_ equals the previously measured ^35^ EC_50_, *b* = 2.5. Therefore, *D*_*eff*_ equals 0.13 μm^2^ / ms. The average diffusion time *t* is, therefore, *t = x*^*2*^*/2D*_*eff*_ *=* 6 min, in rough agreement with the observations. This estimate should be considered approximate in view of the uncertain parameters; nonetheless, diffusion within the hydrogel seems to dominate the waveform of the fluorescence responses.

### Minimal fluorescence from the hydrogel

In addition to advantages of the hydrogel strategy stated in the Introduction, the optical isolation from surrounding tissues and proteins leads to the hope that tissue contributions to F_0_ will be minimal. In measurements not shown, we found that hydrogel fluorescence without introduced iNicSnFR12 was roughly 1/10 the F_0_ value of a hydrogel cast with 10 μM iNicSnFR12. This may allow absolute calibration of *[nicotine]*_*t*_ from measurements of ΔF/F_0_.

### Measurements after storage

To test the rate of biosensor release from the hydrogel, we incubated freshly crosslinked iNicSnFR12 PEGDA/Irgacure hydrogel in 50 mL of 1X PBS, pH 7.4, for three days while shaking. After the incubation, we concentrated the 50 mL of 1X PBS, pH 7.4 to ∼300 μL and we detected no significant fluorescence above background (Data not shown).

As an extreme test of durability, we stored 250 μm slices in 1X PBS, pH7.4 at 4° C for ∼10 months (312 d). Values of F_0_ were 2-3 fold lower than for the fresh hydrogel, and nicotine sensitivity (ΔF/F_0_) was ∼ 2-fold lower than one day after slicing (Figure 2D). We performed further experiments on the storage solution with a fluorescence plate reader. The F_0_ measurements indicated that ∼ 50% of the iNicSnFR12 had diffused from the gel to the storage fluid. In further measurement on the iNicSnFR in the storage fluid, we assessed function by measuring fluorescence excitation spectra ^11^. The ratio between fluorescence excited at 405 nm and 485 nM was ∼1 for fresh solutions of iNicSnFR12 and ∼ 0.75 for the samples stored at 4° C, also suggesting a modest decrease in iNicSnFR12 function during 10 months of storage (Supporting Information: Excitation scans of iNicSnFR12 and 250 um iNicSnFR12 PEGDA hydrogel storage fluid). However, the research uses stated in the Introduction may require only 1-2 days per subject.

### Measurements with an integrated miniature fluorescence microscope

We tested whether the iNicSnFR12 PEGDA/Irgacure hydrogel could be measured with the v4.4 version of the UCLA integrated miniaturized fluorescence microscope **(Figure 3**). The test rig operated the miniscope as an inverted fiber photometer while an iNicSnFR12 PEGDA/Irgacure hydrogel slice was perfused with various [nicotine]. We observed dose-dependent nicotine-induced fluorescence at [nicotine] as low as 10 nM. The 10 nM responses may be distorted by slight movements of the hydrogel.

The miniscope plus fiber optic costs ∼ $2500 US as a kit, roughly 1% of the original cost of the fluorescent microscope that provided the data in Figures 1 and 2. The miniscope provided additional, inexpensive, off-the-shelf proof of concept for the methods. However, its cellphone camera has poorly characterized linearity of responses, which vitiates systematic analysis of the concentration dependence; and as designed, its image rate cannot be slowed below the wasteful 10 Hz. Also, to avoid obvious artifacts from overheating, we decreased the LED power to 20% of maximum. Importantly, the miniscope’s LED, excitation filter, dichroic mirror, and emission filter use the same epifluorescence technology as the fluorescent microscope; and these should be retained in a future, more appropriate, and even less costly specialized miniature fiber photometer.

### Biofluid interactions with iNicSnFR12

In previous reports using fluorescent plate readers, iNicSnFR12 and its predecessors responded to acetylcholine, varenicline, and choline (the latter 100-fold less strongly). ^35^ We explained how these low-MW compounds are unlikely to interfere with *[nicotine]*_*t*_ measurements. ^35^

Dermal ISF cannot yet be isolated in sufficient quantities to serve in a systematic characterization of iNicSnFR12 hydrogels ^42^; therefore, we continued experiments with fluorescent plate readers. Previously, the presence of 25% human serum itself increased background fluorescence ^35^, which we termed F_0’_’. Human serum also apparently contained then-unidentified compound(s) that activated iNicSnFR12. ^35^ With added human serum, the lower limit of quantification increased by 2-3 fold. It was previously not possible to distinguish whether this altered sensitivity arose primarily from the increased F_0_’ or from the background activation of iNicSnFR12 by the endogenous compound(s).

To probe these interactions further, we have tested individual components of human biofluids. Because we evolved the binding moiety of all iDrugSnFRs including iNicSnFR from a choline/betaine-binding binding, and based on our experience with ligands for this series, ^11-15, 44^, we selected 9 additional biogenic amines or alkaloids present at > 1 µM in CSF, ISF, or plasma ^3, 44^ as candidates to activate iNicSnFR12. ISF concentrations are only partially characterized ^45^; therefore, we cite plasma concentrations ^3, 44^ in µM: tryptamine (16.5), putrescine (8.4),, cadaverine (2.0), tyramine (1.8), spermidine (3.1), urea 4.0, L-tryptophan (65), sarcosine (1.4), and carnitine (50). Of these, only carnitine consistently gave a detectable ΔF/F_0_ at the plasma concentration (Supporting information: Concentration-response relations for biogenic amines and alkaloids with iNicSnF12). Referring these fluorescent plate reader data to comparable previous data ^35^, this carnitine response would equal the response to ∼ 15 nM nicotine, well within the linear range and unlikely to affect sensitivity by itself. Nonetheless, we are considering protective layers that might exclude carnitine. On the one hand, Nafion™, an exemplar tetrafluoroethylene based fluoropolymer copolymer, excludes nicotine and would not serve as a protective layer ^33^. On the other hand, PDMS is quite permeant to nicotine ^46, 47^ and would likely exclude carnitine, choline, and acetylcholine--all quaternary amines.

The minimal effects of endogenous biogenic amines and alkaloids imply that the more important challenge is to suppress the increased F_0_’ of biofluids. We previously suggested that the elevation arises mostly from bilirubin ^35^. Because bilirubin is usually conjugated to albumin, ^43^ we hypothesize that the hydrogel tactic will decrease F_0_’ by excluding albumin. A decisive test of this hypothesis will require redesigning the optics to record fluorescence from only the hydrogel, rather than including surrounding test fluids or tissues. In summary, it is likely that a combination of the hydrogel strategy and protective layers can optimize an iNicSnFR12-based continuous nicotine monitor.

## Conclusions

An engineered, bacterially expressed, purified, fluorescent nicotine biosensor protein can be encapsulated in several hydrogel formulations, while retaining the ability to continuously detect dose-dependent signals over several log units of [nicotine]. The most successful iteration, utilizing PEGDA/Irgacure, was readily sliced by a vibratome, and showed minimal leaching of iNicSnFR protein on a time scale of days after preparation.

After nicotine appears at the hydrogel surface, diffusion slows access to the iNicSnFR molecules. We explain how diffusion is slowed slightly by the gel and markedly by rebinding to iNicSnFR12 itself. But iNicSnFR12 must be concentrated sufficiently to give a good signal-to-noise (S/N) ratio. This three-effect optimization--buffering, diffusion, S/N—also appears importantly in the neuroscience literature on genetically encoded fluorescent Ca^2+^ sensor proteins. For measurements that resolve the ∼ 5 min bolus of nicotine from smoking or vaping, we conclude that iNicSnFR molecules must be present at 10 µM and within ∼ 200 μm of the biofluid surface.

The approximately linear dependence of induced fluorescence on [nicotine] and the minimal fluorescence of the gel itself lead to the conclusion that absolute calibration of ΔF/F_0_ in terms of *[nicotine]*_*t*_ may be possible in tissue. We conclude that, given additional resources, a minimally invasive fiber-optic based continuous monitor measuring *[nicotine]*_*t*_ in interstitial fluid can eventually be developed with a form factor and cost comparable to a modern amperometric continuous glucose monitor, for research on persons who ingest nicotine. Further engineering ^48^ would improve the sensitivity of the iNicSnFR series, further integrate the electronics, optimize the optics including light shielding, adhere or attach the hydrogel to the fiber optic, test protective coatings, and develop an insertion device.

Finally, although we did not test the immobilization of previously reported biosensors for other drug classes (iDrugSnFRs),^11-14, 16, 44^ photophysics and stability among iDrugSnFRs is nearly indistinguishable, which should result in comparable results if they were similarly encased. This would expand the range of therapeutic and abused drugs that could be monitored continuously in a hydrogel.

## ACKNOWLEDGEMENTS

We thank Neal Benowitz, Bruce Cohen, Mario Danek, Sophie Dalfonso, Sujit Datta, Ryan Drenan, Nick Friesenhahn, Heather Lukas, Anand Muthusamy, and Koji Sode for advice. We thank Carlos Lois for use of a vibratome.

Funding was provided by the Caltech Merkin Insitute for Translational Research, the Caltech Sensing to Innovation (S2I) Fund, the Caltech Rosen Bioengineering Center, the Caltech Rothenberg Innovation Initiative (RI^2^), and the Caltech Carver Mead New Ventures Fund.

## REFERENCES

1. Benowitz, N. L., Pharmacokinetic considerations in understanding nicotine dependence. Ciba Found Symp 1990, 152, 186–200; discussion 200-9.

2. Benowitz, N. L.; Jacob, P., 3rd; Denaro, C.; Jenkins, R., Stable isotope studies of nicotine kinetics and bioavailability. Clin Pharmacol Ther 1991, 49 (3), 270–7.

3. Matta, S. G.; Balfour, D. J.; Benowitz, N. L.; Boyd, R. T.; Buccafusco, J. J.; Caggiula, A. R.; Craig, C. R.; Collins, A. C.; Damaj, M. I.; Donny, E. C.; Gardiner, P. S.; Grady, S. R.; Heberlein, U.; Leonard, S. S.; Levin, E. D.; Lukas, R. J.; Markou, A.; Marks, M. J.; McCallum, S. E.; Parameswaran, N.; Perkins, K. A.; Picciotto, M. R.; Quik, M.; Rose, J. E.; Rothenfluh, A.; Schafer, W. R.; Stolerman, I. P.; Tyndale, R. F.; Wehner, J. M.; Zirger, J. M., Guidelines on nicotine dose selection for in vivo research. Psychopharm 2007, 190 (3), 269–319.

4. Kovar, L.; Selzer, D.; Britz, H.; Benowitz, N.; St Helen, G.; Kohl, Y.; Bals, R.; Lehr, T., Comprehensive Parent-Metabolite PBPK/PD Modeling Insights into Nicotine Replacement Therapy Strategies. Clin Pharmacokinet 2020, 59 (9), 1119–1134.

5. Lunell, E.; Fagerstrom, K.; Hughes, J.; Pendrill, R., Pharmacokinetic Comparison of a Novel Non-tobacco-Based Nicotine Pouch (ZYN) With Conventional, Tobacco-Based Swedish Snus and American Moist Snuff. Nicotine Tob Res 2020, 22 (10), 1757–1763.

6. Unger, E. K.; Keller, J. P.; Altermatt, M.; Liang, R.; Matsui, A.; Dong, C.; Hon, O. J.; Yao, Z.; Sun, J.; Banala, S.; Flanigan, M. E.; Jaffe, D. A.; Hartanto, S.; Carlen, J.; Mizuno, G. O.; Borden, P. M.; Shivange, A. V.; Cameron, L. P.; Sinning, S.; Underhill, S. M.; Olson, D. E.; Amara, S. G.; Temple Lang, D.; Rudnick, G.; Marvin, J. S.; Lavis, L. D.; Lester, H. A.; Alvarez, V. A.; Fisher, A. J.; Prescher, J. A.; Kash, T. L.; Yarov-Yarovoy, V.; Gradinaru, V.; Looger, L. L.; Tian, L., Directed Evolution of a Selective and Sensitive Serotonin Sensor via Machine Learning. Cell 2020, 183 (7), 1986-2002.e26.

7. Marvin, J. S.; Shimoda, Y.; Magloire, V.; Leite, M.; Kawashima, T.; Jensen, T. P.; Kolb, I.; Knott, E. L.; Novak, O.; Podgorski, K.; Leidenheimer, N. J.; Rusakov, D. A.; Ahrens, M. B.; Kullmann, D. M.; Looger, L. L., A genetically encoded fluorescent sensor for in vivo imaging of GABA. Nat Methods 2019, 16 (8), 763–770.

8. Marvin, J. S.; Borghuis, B. G.; Tian, L.; Cichon, J.; Harnett, M. T.; Akerboom, J.; Gordus, A.; Renninger, S. L.; Chen, T. W.; Bargmann, C. I.; Orger, M. B.; Schreiter, E. R.; Demb, J. B.; Gan, W. B.; Hires, S. A.; Looger, L. L., An optimized fluorescent probe for visualizing glutamate neurotransmission. Nat Methods 2013, 10 (2), 162–70.

9. Patriarchi, T.; Cho, J. R.; Merten, K.; Howe, M. W.; Marley, A.; Xiong, W. H.; Folk, R. W.; Broussard, G. J.; Liang, R.; Jang, M. J.; Zhong, H.; Dombeck, D.; von Zastrow, M.; Nimmerjahn, A.; Gradinaru, V.; Williams, J. T.; Tian, L., Ultrafast neuronal imaging of dopamine dynamics with designed genetically encoded sensors. Science 2018, 360 (6396), eaat4422.

10. Robinson, J. E.; Coughlin, G. M.; Hori, A. M.; Cho, J. R.; Mackey, E. D.; Turan, Z.; Patriarchi, T.; Tian, L.; Gradinaru, V., Optical dopamine monitoring with dLight1 reveals mesolimbic phenotypes in a mouse model of neurofibromatosis type 1. Elife 2019, 8.

11. Bera, K.; Kamajaya, A.; Shivange, A. V.; Muthusamy, A. K.; Nichols, A. L.; Borden, P. M.; Grant, S.; Jeon, J.; Lin, E.; Bishara, I.; Chin, T. M.; Cohen, B. N.; Kim, C. H.; Unger, E. K.; Tian, L.; Marvin, J. S.; Looger, L. L.; Lester, H. A., Biosensors Show the Pharmacokinetics of S-Ketamine in the Endoplasmic Reticulum. Frontiers in cellular neuroscience 2019, 13, 499–499.

12. Muthusamy, A. K.; Kim, C. H.; Virgil, S. C.; Knox, H. J.; Marvin, J. S.; Nichols, A. L.; Cohen, B. N.; Dougherty, D. A.; Looger, L. L.; Lester, H. A., Three Mutations Convert the Selectivity of a Protein Sensor from Nicotinic Agonists to S-Methadone for Use in Cells, Organelles, and Biofluids. J Am Chem Soc 2022, 144 (19), 8480–8486.

13. Nichols, A. L.; Blumenfeld, Z.; Fan, C.; Luebbert, L.; Blom, A. E. M.; Cohen, B. N.; Marvin, J. S.; Borden, P. M.; Kim, C.; Muthusamy, A. K.; Shivange, A. V.; Knox, H. J.; Campello, H. R.; Wang, J. H.; Dougherty, D. A.; Looger, L. L.; Gallagher, T.; Rees, D. C.; Lester, H. A., Fluorescence Activation Mechanism and Imaging of Drug Permeation with New Sensors for Smoking-Cessation Ligands. eLife 2022, 11, e74648.

14. Nichols, A. L.; Blumenfeld, Z.; Luebbert, L.; Knox, H. J.; Muthusamy, A. K.; Marvin, J. S.; Kim, C. H.; Grant, S. N.; Walton, D. P.; Cohen, B. N.; Hammar, R.; Looger, L.; Artursson, P.; Dougherty, D. A.; Lester, H. A., Selective Serotonin Reuptake Inhibitors within Cells: Temporal Resolution in Cytoplasm, Endoplasmic Reticulum, and Membrane. J Neurosci 2023, 43, 2222–41.

15. Shivange, A. V.; Borden, P. M.; Muthusamy, A. K.; Nichols, A. L.; Bera, K.; Bao, H.; Bishara, I.; Jeon, J.; Mulcahy, M. J.; Cohen, B.; O′Riordan, S. L.; Kim, C.; Dougherty, D. A.; Chapman, E. R.; Marvin, J. S.; Looger, L. L.; Lester, H. A., Determining the pharmacokinetics of nicotinic drugs in the endoplasmic reticulum using biosensors. J Gen Physiol 2019, 151 (4), 738–757.

16. Beatty, Z. G.; Muthusamy, A. K.; Unger, E. K.; Dougherty, D. A.; Tian, L.; Looger, L. L.; Shivange, A. V.; Bera, K.; Lester, H. A.; Nichols, A. L., Fluorescence Screens for Identifying Central Nervous System-Acting Drug-Biosensor Pairs for Subcellular and Supracellular Pharmacokinetics. Bio Protoc 2022, 12 (22).

17. El-Sherbiny, I. M.; Yacoub, M. H., Hydrogel scaffolds for tissue engineering: Progress and challenges. Glob Cardiol Sci Pract 2013, 2013 (3), 316–42.

18. Gao, Y.; Zhang, X.; Zhou, H., Biomimetic Hydrogel Applications and Challenges in Bone, Cartilage, and Nerve Repair. Pharmaceutics 2023, 15 (10).

19. Mortellaro, M.; DeHennis, A., Performance characterization of an abiotic and fluorescent-based continuous glucose monitoring system in patients with type 1 diabetes. Biosens Bioelectron 2014, 61, 227–31.

20. Lavrentev, F. V.; Shilovskikh, V. V.; Alabusheva, V. S.; Yurova, V. Y.; Nikitina, A. A.; Ulasevich, S. A.; Skorb, E. V., Diffusion-Limited Processes in Hydrogels with Chosen Applications from Drug Delivery to Electronic Components. Molecules 2023, 28 (15).

21. Schmieg, B.; Dobber, J.; Kirschhofer, F.; Pohl, M.; Franzreb, M., Advantages of Hydrogel-Based 3D-Printed Enzyme Reactors and Their Limitations for Biocatalysis. Front Bioeng Biotechnol 2018, 6, 211.

22. Rollema, H.; Shrikhande, A.; Ward, K. M.; Tingley, F. D., 3rd; Coe, J.W.; O′Neill, B. T.; Tseng, E.; Wang, E. Q.; Mather, R. J.; Hurst, R. S.; Williams, K. E.; de Vries, M.; Cremers, T.; Bertrand, S.; Bertrand, D., Pre-clinical properties of the α4β2 nicotinic acetylcholine receptor partial agonists varenicline, cytisine and dianicline translate to clinical efficacy for nicotine dependence. Br J Pharmacol 2010, 160 (2), 334–45.

23. Dempsey, D.; Tutka, P.; Jacob, P., 3rd; Allen, F.; Schoedel, K.; Tyndale, R. F.; Benowitz, N. L., Nicotine metabolite ratio as an index of cytochrome P450 2A6 metabolic activity. Clin Pharmacol Ther 2004, 76 (1), 64–72.

24. Henderson, B. J.; Lester, H. A., Inside-out neuropharmacology of nicotinic drugs. Neuropharmacol 2015, 96 (Pt B), 178–93.

25. Lerman, C.; Schnoll, R. A.; Hawk, L. W., Jr.; Cinciripini, P.; George, T. P.; Wileyto, E. P.; Swan, G. E.; Benowitz, N. L.; Heitjan, D. F.; Tyndale, R. F.; Group, P.-P. R., Use of the nicotine metabolite ratio as a genetically informed biomarker of response to nicotine patch or varenicline for smoking cessation: a randomised, double-blind placebo-controlled trial. Lancet Respir Med 2015, 3 (2), 131–138.

26. Chenoweth, M. J.; Tyndale, R. F., Pharmacogenetic Optimization of Smoking Cessation Treatment. Trends Pharmacol Sci 2017, 38 (1), 55–66.

27. Benowitz, N. L.; Jacob, P., 3rd; Fong, I.; Gupta, S., Nicotine metabolic profile in man: comparison of cigarette smoking and transdermal nicotine. J Pharmacol Exp Ther 1994, 268 (1), 296–303.

28. Solingapuram Sai, K. K.; Zuo, Y.; Rose, J. E.; Garg, P. K.; Garg, S.; Nazih, R.; Mintz, A.; Mukhin, A. G., Rapid Brain Nicotine Uptake from Electronic Cigarettes. J Nucl Med 2020, 61 (6), 928–930.

29. Chenoweth, M. J.; Lerman, C.; Knight, J.; Tyndale, R. F., Influence of CYP2A6 Genetic Variation, Nicotine Dependence Severity, and Treatment on Smoking Cessation Success. Nicotine Tob Res 2023, 25 (6), 1207–1211.

30. El-Boraie, A.; Taghavi, T.; Chenoweth, M. J.; Fukunaga, K.; Mushiroda, T.; Kubo, M.; Lerman, C.; Nollen, N. L.; Benowitz, N. L.; Tyndale, R. F., Evaluation of a weighted genetic risk score for the prediction of biomarkers of CYP2A6 activity. Addict Biol 2020, 25 (1), e12741.

31. Mehmeti, E.; Kilic, T.; Laur, C.; Carrara, S., Electrochemical determination of nicotine in smokers’ sweat. Microchemical Journal 2020, 158.

32. Tai, L. C.; Ahn, C. H.; Nyein, H. Y. Y.; Ji, W.; Bariya, M.; Lin, Y.; Li, L.; Javey, A., Nicotine Monitoring with a Wearable Sweat Band. ACS sensors 2020, 5 (6), 1831–1837.

33. Galagan, J.; Grinstaff, M.; Kuzmanovic, U.; Chen, M.; Alexandrovna, M.; Allen, K. Enzyme-Based Electrochemical Nicotine Biosensor. 11,331,020, 2022.

34. Min, J.; Tu, J.; Xu, C.; Lukas, H.; Shin, S.; Yang, Y.; Solomon, S. A.; Mukasa, D.; Gao, W., Skin-Interfaced Wearable Sweat Sensors for Precision Medicine. Chem Rev 2023, 123 (8), 5049–5138.

35. Haloi, N.; Huang, S.; Nichols, A. L.; Fine, E. J.; Friesenhahn, N. J.; Marotta, C. B.; Dougherty, D. A.; Lindahl, E.; Howard, R. J.; Mayo, S. L.; Lester, H. A., Interactive computational and experimental approaches improve the sensitivity of periplasmic binding protein-based nicotine biosensors for measurements in biofluids. Protein Eng Des Sel 2024, 37.

36. Srinivasan, R.; Pantoja, R.; Moss, F. J.; Mackey, E. D. W.; Son, C.; Miwa, J.; Lester, H.A., Nicotine upregulates α4β2 nicotinic receptors and ER exit sites via stoichiometry-dependent chaperoning. J. Gen .Physiol. 2011, 137, 59–79.

37. Shivange, A. V.; Borden, P. M.; Muthusamy, A. K.; Nichols, A. L.; Bera, K.; Bao, H.; Bishara, I.; Jeon, J.; Mulcahy, M. J.; Cohen, B.; O′Riordan, S. L.; Kim, C.; Dougherty, D. A.; Chapman, E. R.; Marvin, J. S.; Looger, L. L.; Lester, H. A., Determining the pharmacokinetics of nicotinic drugs in the endoplasmic reticulum using biosensors. J Gen Physiol 2019, 151 (6), 738–757.

38. Dong, Z.; Mau, W.; Feng, Y.; Pennington, Z. T.; Chen, L.; Zaki, Y.; Rajan, K.; Shuman, T.; Aharoni, D.; Cai, D. J., Minian, an open-source miniscope analysis pipeline. Elife 2022, 11.

39. Ghosh, K. K.; Burns, L. D.; Cocker, E. D.; Nimmerjahn, A.; Ziv, Y.; Gamal, A. E.; Schnitzer, M. J., Miniaturized integration of a fluorescence microscope. Nat Methods 2011, 8 (10), 871–8.

40. Crank, J., The Mathematics of Diffusion. Second ed.; Clarendon Press: Oxford, 1975.

41. Wathey, J. C.; Nass, M. N.; Lester, H. A., Numerical reconstruction of the quantal event at nicotinic synapses. Biophys. J. 1979, 27, 145–164.

42. Friedel, M.; Thompson, I. A. P.; Kasting, G.; Polsky, R.; Cunningham, D.; Soh, H. T.; Heikenfeld, J., Opportunities and challenges in the diagnostic utility of dermal interstitial fluid. Nat Biomed Eng 2023, 7 (12), 1541–1555.

43. Wolfbeis, O. S.; Leiner, M., MAPPING OF THE TOTAL FLUORESCENCE OF HUMAN-BLOOD SERUM AS A NEW METHOD FOR ITS CHARACTERIZATION. ANALYTICA CHIMICA ACTA 1985, 167 (JAN), 203–215.

44. Blumenfeld, Z., Genetically Encoded Biosensors for Ketamine and Other Rapidly Acting Antidepressants in Zebrafish and Cell Culture. Ph D. Thesis, California Institute of Technology: 2023.

45. Kolluru, C.; Williams, M.; Yeh, J. S.; Noel, R. K.; Knaack, J.; Prausnitz, M. R., Monitoring drug pharmacokinetics and immunologic biomarkers in dermal interstitial fluid using a microneedle patch. Biomed Microdevices 2019, 21 (1), 14.

46. Baltussen, E.; den Boer, A.; Sandra, P.; Janssen, H. G.; Cramers, C., Monitoring of nicotine in air using sorptive enrichment on polydimethylsiloxane and TD-CGC-NPD. Chromatographia 1999, 49 (9), 520–524.

47. Godage, N. H.; Cudjoe, E.; Neupane, R.; Boddu, S. H.; Bolla, P. K.; Renukuntla, J.; Gionfriddo, E., Biocompatible SPME fibers for direct monitoring of nicotine and its metabolites at ultra trace concentration in rabbit plasma following the application of smoking cessation formulations. J Chromatogr A 2020, 1626, 461333.

48. Clark, H. A., Has Sensing Become an Engineering Discipline? ACS sensors 2020, 5 (2), 292–293.

