## Supporting Information for "Hydrogel encapsulation of a designed fluorescent protein biosensor for continuous measurements of sub-100 nanomolar nicotine"

**Nichols AL, Marotta CB, et al. 2024 ACS Sensors Supplemental Information**

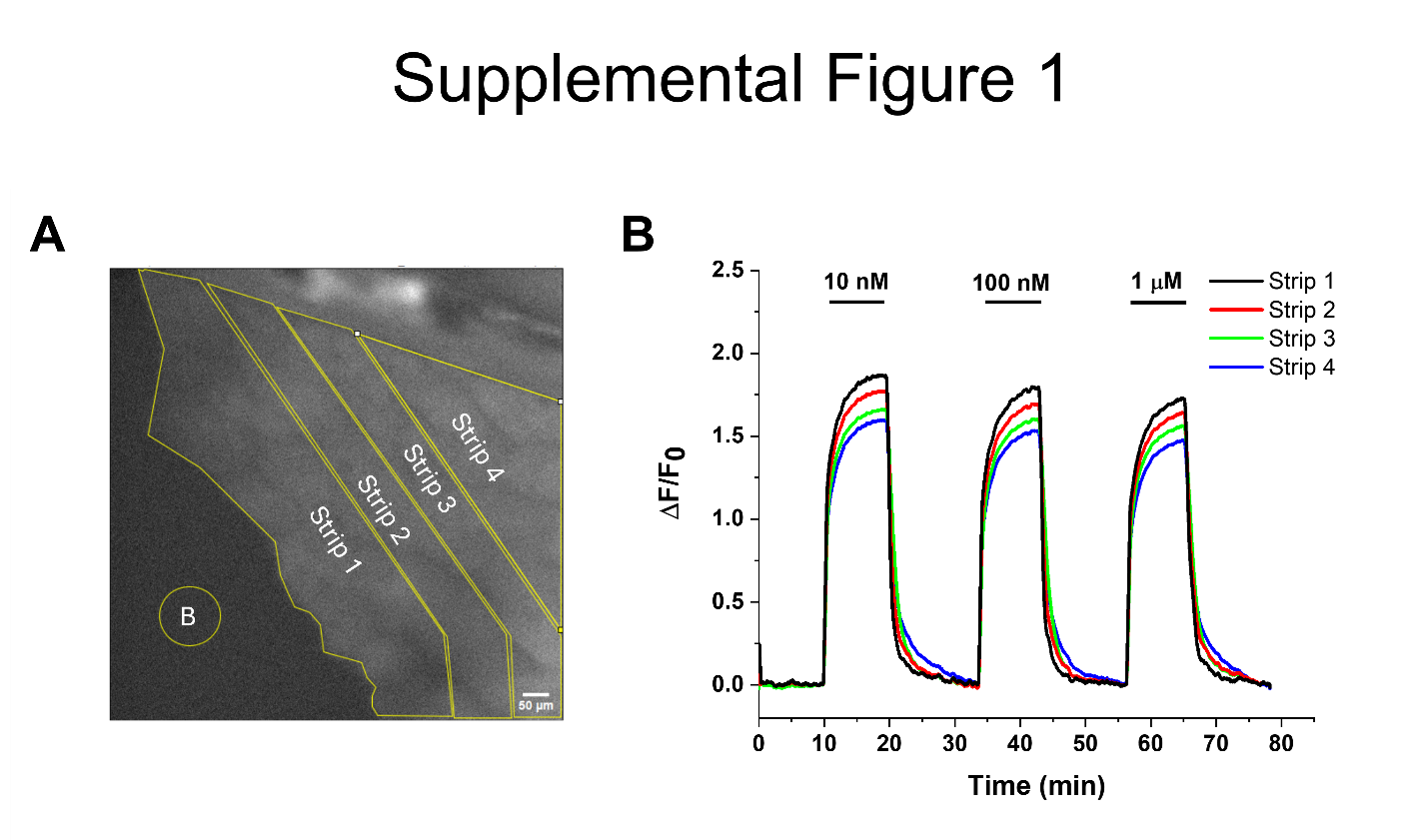

**Analysis of solvent proximity to fluorescent response of iNicSnFR12 acrylamide hydrogel.** (A) Division of an iNicSnFR12 acrylamide hydrogel into different strips for analysis. The encircled B indicates the background used in calculations. (B) Sections of acrylamide hydrogel located more distally from a drug bath have slower wash in and wash-out kinetics in comparison to those more proximally located.

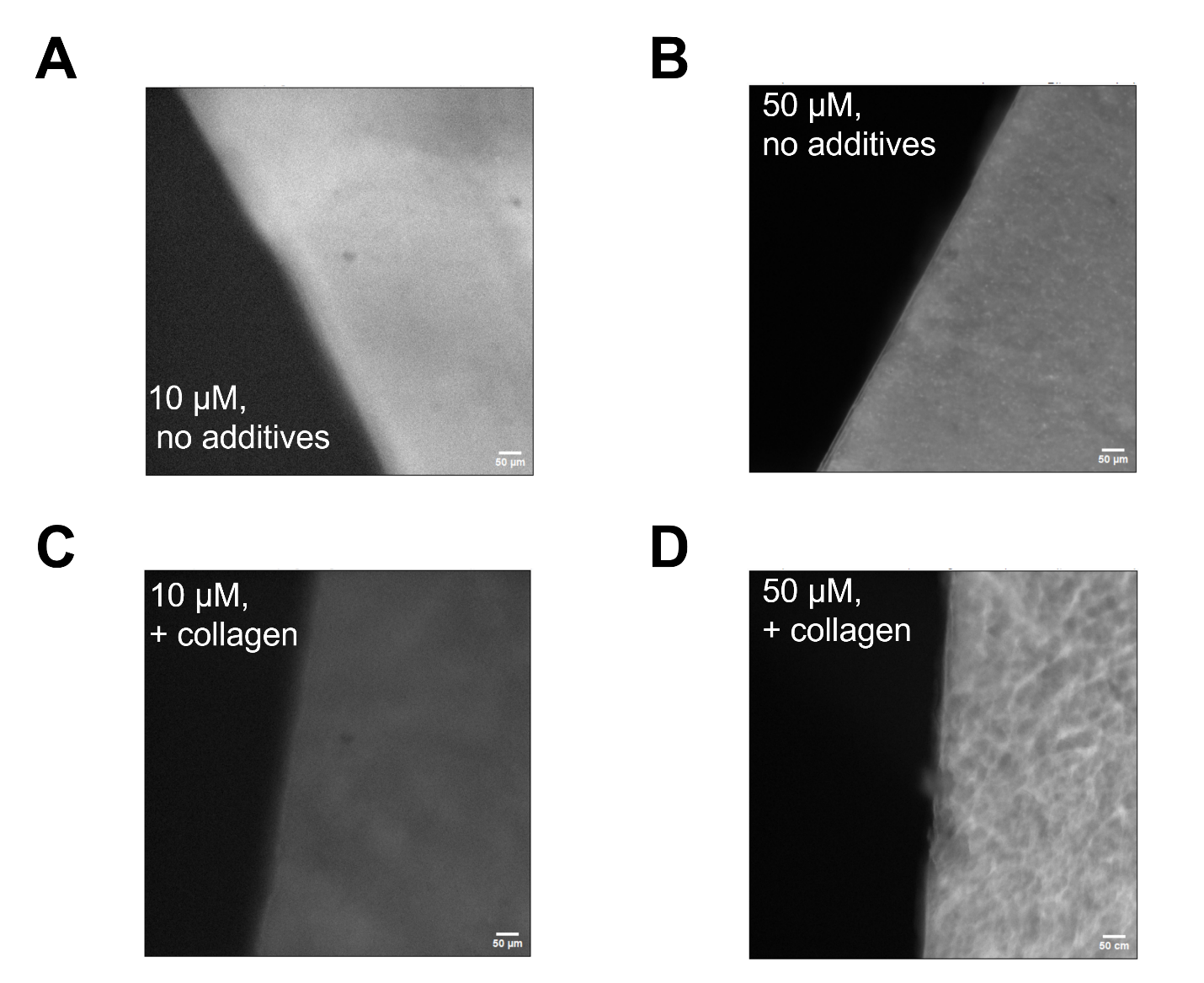

**Widefield imaging of various PEGDA hydrogel preparations.** (A) A 10 μM iNicSnFR12 PEGDA/Irgacure has a homogeneous distribution while a similar hydrogel (B) with 50 μM iNicSnFR12 contains puncta. (C) When collagen is included in a 10 μM iNicSnFR12 PEGDA/Irgacure, a banded pattern is observed that is markedly increased (D) when the iNicSnFR12 concentration is increased to 50 μM.

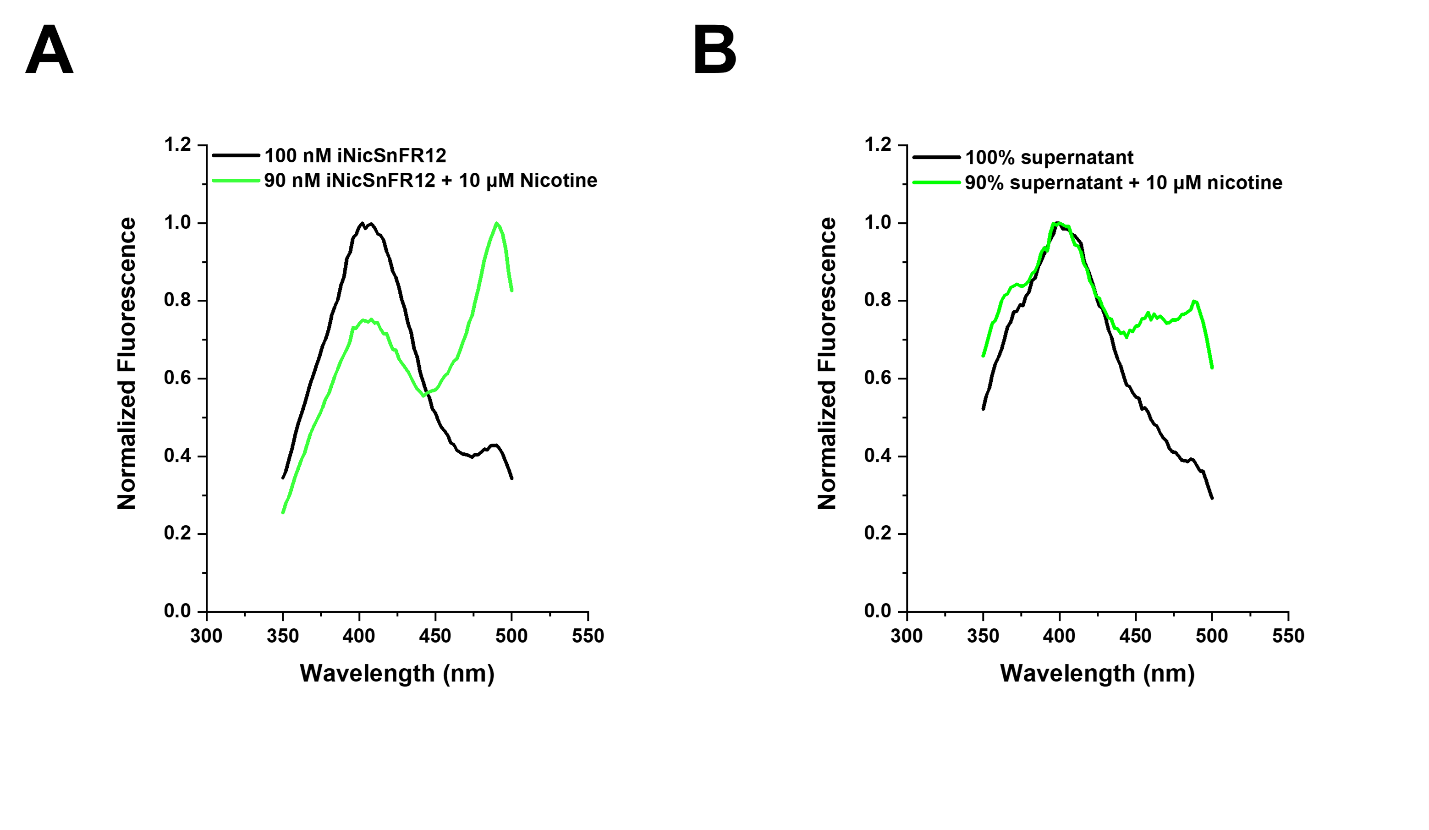

**Excitation scans of iNicSnFR12 and 250 um iNicSnFR12 PEG-DA hydrogel storage fluid.** (A) iNicSnFR12 contains the expected peaks expected from the absorption peaks of iNicSnFR12 (ref. 34) (405 nm, 496 nm). Addition of nicotine decreases the 405 nm peak slightly more than the 10% expected from dilution; and the 496 nm peak increases dramatically. (B) Supernatant from a 10-month old hydrogel preparation contains a peak at 405 nm, but this is not decreased by the addition of nicotine. Addition of nicotine increases the 496 nm peak less dramatically

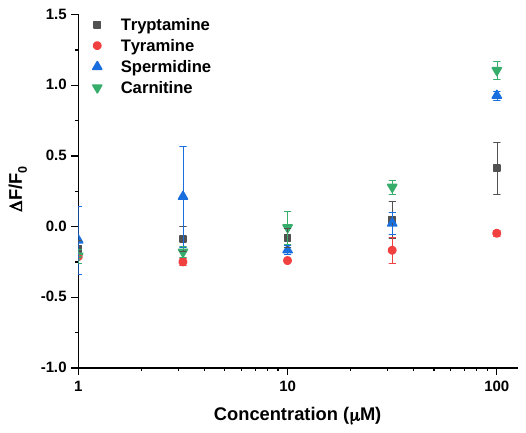

**Concentration-response relations for biogenic amines and alkaloids with iNicSnF12**.

Tests were conducted in triplicate with 100 nM iNicSnFR12 in a Tecan Spark M20 plate reader, 1X PBS, pH 7.4. Putrescince, cadaverine, urea, and L-tryptophan, and sarcosine yielded ΔF/F_0_ < 0.1 at plasma levels and are not shown. Concentration-response relations were performed for tryptamine, tyramine, spermidine, and carnitine in triplicate. Mean ± SD are shown.
